# Advanced Age Has Dissociable Effects on Hippocampal CA1 and CA3 Ripples

**DOI:** 10.1101/2021.08.27.457373

**Authors:** Nicholas M. DiCola, Alexa L. Lacy, Omar J. Bishr, Kathryn M. Kimsey, Jenna L. Whitney, Sarah D. Lovett, Sara N. Burke, Andrew P. Maurer

## Abstract

Sharp-wave/ripples are brief, high-frequency events in hippocampal subregions CA3 and CA1 that occur during rest or pauses in behavior. Ripples detected in CA1 have lower frequency in aged compared to young rats. Although CA1 ripples are theorized to manifest from CA3, ripple dynamics in CA3 have not been examined in aged animals. The current study obtained simultaneous recordings between CA1 and CA3 in young and aged rats to examine sharp-wave/ripple characteristics in both regions in relation to age. While CA1 ripple frequency was reduced with age, there were no age differences in CA3 ripples. In aged, but not young, rats there was also a significant increase in the probability of CA3 and CA1 ripples co-occurring between the pre- and post-behavior rest epochs. Moreover, in both age groups, CA1 ripples that co-occurred with a CA3 ripple had increased frequency, power, and duration. These findings suggest age differences in CA1 are not due to altered afferent input from CA3, but instead reflect distinct mechanisms of ripple generation with age.

**HIGHLIGHTS:**

- CA1 ripple frequency is reduced with age.
- CA3 ripple characteristics do not change with age.
- In aged rats only, CA3-CA1 ripple co-occurrence increased following behavior.
- CA1 ripples that co-occurred with CA3 had greater frequency, power, and duration.

## 1 INTRODUCTION

As the world’s demographics shift towards proportionally more older adults than ever seen before in recorded history, age-related cognitive decline that threatens quality of life is a growing world-wide problem (Blazer et al., 2015). The average life expectancy has outpaced the cognitive health span, with a majority of adults over the age of 65 suffering from a variety of decreased cognitive abilities. One brain region that is well documented as being particularly vulnerable to neurobiological alterations in advanced age is the hippocampus. As such, performance on behavioral tasks that rely on the hippocampus, such as explicit memory (Rosenzweig and Barnes, 2003) and spatial navigation (Lester et al., 2017), are degraded in aged humans (Blazer et al., 2015), monkeys (Rapp et al., 1997), and rodents (Barnes, 1979). Our understanding of the biological mechanisms that account for these deficits, however, is incomplete.

Normative aging is not associated with profound cell loss (West et al., 1994; Rapp and Gallagher, 1996), suggesting that the mechanisms of impaired hippocampal function in advanced age must arise from more subtle neurobiological changes that affect synaptic function and neuron signaling to alter the dynamics of neuronal network activity that support behavior (Rosenzweig and Barnes, 2003; Burke and Barnes, 2006). For instance, the evoked field excitatory post-synaptic potential (fEPSP) at the CA3 to CA1 Schaffer collateral synapse is reduced in aged compared to young rats (Landfield et al., 1986; Barnes et al., 1992; Deupree et al., 1993; Tombaugh et al., 2002). As the pre-synaptic fiber potential of the Schaffer Collaterals does not change between young and aged animals (Barnes et al., 1997; Rosenzweig et al., 1997; Barnes et al., 2000; Tombaugh et al., 2002), it suggests that the number of afferent projections are intact. Therefore, decreased fEPSP in advanced age is likely related to the decline in postsynaptic densities of perforated synapses in CA1 (Nicholson et al., 2004) and impaired temporal summation in CA1 pyramidal neurons (Rosenzweig et al., 1997). Finally, there are notable age-related changes in the activity properties of CA3 neurons that are likely to impact efferent targets in CA1. These changes include a decrease in interneurons expressing GAD67 (Spiegel et al., 2013), and elevated activity in CA3 excitatory cells (Wilson et al., 2005b; Robitsek et al., 2015; Thomé et al., 2016; Lee et al., 2021). Together, these data suggest that the ability of the CA3 subregion to influence CA1 activity is likely to be altered in advanced age. To our knowledge, however, there have been no investigations regarding whether or not coordinated activity between CA3 and CA1 is altered as a function of age in awake behaving animals.

A readout of coordinated synaptic activity are oscillations in the local field potential (LFP), which are the superposition of inhibitory and excitatory synaptic events (Buzsáki et al., 1983). Oscillations in the high frequency range (>120 Hz) (Ylinen et al., 1995), known as ripples, are thought to be the product of mass CA3 depolarization of CA1 pyramidal neurons at the Schaffer collateral synapse, known as the sharp wave, which occur during slow-wave sleep (Wilson and McNaughton, 1994; Chrobak and Buzsaki, 1996; Skaggs and McNaughton, 1996; Kudrimoti et al., 1999; Nadasdy et al., 1999; Sutherland and McNaughton, 2000; Lee and Wilson, 2002; Buzsáki et al., 2003; Born et al., 2006; Grosmark et al., 2012; Cox et al., 2020; Laventure and Benchenane, 2020; Ngo et al., 2020; Oyanedel et al., 2020; Skelin et al., 2021) or periods of quiet wakefulness (Buzsaki, 1986; Chrobak and Buzsaki, 1996; Foster and Wilson, 2006; Jackson et al., 2006; O’Neill et al., 2006; Csicsvari et al., 2007; Diba and Buzsáki, 2007; Davidson et al., 2009; Karlsson and Frank, 2009; Sullivan et al., 2011; Buzsáki, 2015; Giri et al., 2019). Although ripples in CA1 are most evident in the pyramidal cell body layer, they originate from mass depolarization of the apical dendrites in the stratum radiatum, plausibly driven by CA3 input (Ylinen et al., 1995). Therefore, these events could be a neurophysiological indicator of the integrity of CA3 to CA1 coordination.

Previous work has described differences in CA1 ripples between young and aged rats (Wiegand et al., 2016; Cowen et al., 2020). In old rats, ripples have a reduced peak frequency compared to young animals and a lowered rate of occurrence after the rats performed a circular running task involving acquisition of an association between a spatial location and eyelid stimulation. These changes in CA1 ripples with age may have implications for cognitive aging, as it has been proposed that ripples support memory function (Chrobak and Buzsaki, 1996; Nakashiba et al., 2009; Buzsáki, 2015). They may also reflect synaptic reorganization that occurs during behavior (Foster and Wilson, 2006; Csicsvari et al., 2007; Davidson et al., 2009; Atherton et al., 2015; Buzsáki, 2015; Ambrose et al., 2016; Deng et al., 2016; Cazin et al., 2020), although some findings have challenged that hypothetical role of reactivation during ripples for memory stabilization (Gupta et al., 2010; Kovács et al., 2016). Critically, while ripples are known to require intact inhibitory and excitatory balance (Stark et al., 2014; Gan et al., 2017) and there are age-related disruption in inhibition within CA3 (Shetty and Turner, 1998; Shi et al., 2004; Stanley and Shetty, 2004) that are presumed to result in increased excitation of pyramidal neurons (Wilson et al., 2005b; Lee et al., 2021), the sharp wave/ripple characteristics in this subregion of the hippocampus have not been investigated in aged rats.

## 2 MATERIALS AND METHODS

### 2.01 Subjects

The current experiments used 8 aged (24-26 month) and 10 young (4-8 month) male, Fischer 344xBrown Norway F1 hybrid rats obtained from the National Institute on Aging (NIA) colony at Charles River. One aged and two young rats were excluded for not having simultaneous CA1 and CA3 electrophysiological recordings for a final total of 7 aged and 8 young rats. While the importance of the inclusion of both sexes is recognized, a lack of availability of female rats from the NIA colony during the timeframe that the current data were collected and then the research closure during the COVID-19 pandemic precluded the use of female rats in this study. Upon arrival to the animal colony at the University of Florida, the animals habituated for one week before beginning food restriction and behavioral shaping. Rats were restricted to 85% of their *ad lib* weight. In some cases, rats were over conditioned upon arrival at the colony and 85% of the initial weight was insufficient to achieve a healthy body condition to motivate appetitive behaviors. This was observed in the all the aged rats that arrive at the facility over conditioned with excessive visceral white adipose depots (Hernandez et al., 2018). Thus, throughout the period of restricted feeding, in addition to daily weighing, rats were assessed weekly for a body condition score ranging from 1-5, with 3 being optimal, 1 being emaciated and 5 being obese. The body condition score is assigned based on the presence of palpable fat deposits over the lumbar vertebrae and pelvic bones (Ullman-Culleré and Foltz, 1999; Hickman and Swan, 2010). Baseline weights were re-set to correspond to the weight of the animal with a body condition score of 3. If rats dropped to a body condition of 2.5 more food was made available. Animals were individually housed and maintained on a reverse 12:12 hour light:dark cycle, with training and neurophysiological recordings occurring during the dark phase. All experiments followed the guidelines of the United States National Institute of Health’s *Guide for the Care and Use of Laboratory Animals* and were approved by the University of Florida Institutional Animal Care and Use Committee.

One week after arriving to the animal housing facility rats began habituation to experimenters and the maze environment. They were placed into the room and trained to traverse their respective maze until they could successfully complete 30 laps in 30 minutes. Cereal marshmallows broken into approximately .03-.1 gram pieces were used as a reward. This was followed by object displacement training where the animals learned to move a 6.25×6.25×5.75 cm square LEGO© block in preparation for object discrimination. After successfully reaching criterion on object displacement training, rats were then trained on one of several different behavioral tasks: the LEGO© discrimination task (Johnson et al., 2017), the Object-Place Paired Association task (Hernandez et al., 2015), or continuous alternation. Not all animals were able to complete all behavioral tasks before the implants were no longer viable or the animal had health complications related to age.

### 2.02 Surgery and Neurophysiology

After reaching criterion, rats were stereotaxically implanted with dual, chronic, high-density silicon probes from Cambridge Neurotech mounted on moveable nano-drives with 5 mm of vertical travel. Three days before surgery, rats were given an anti-biotic regiment that continued until 10 days post-surgery (.5 mL/day of sulfatrim pediatric suspension: 20 mg Sulfamethoxazole, 4 mg Trimethoprim, no more than .5% alcohol by volume). Immediately prior to surgery all rats were also administered .5 mL of Metacam diluted with .1 mL of sterile saline and 0.01 mL/100 grams of bodyweight of Atropine diluted with .1 mL of sterile saline. Rats were initially put under anesthesia with 5% by volume isoflurane that was then lowered to 1-2% throughout surgery. Oxygen flow rate was maintained between 1.5-2.0 L/ minute. Group 1 (4 young, 4 aged) was then implanted with dual F-series silicon probes in the cortex above the hippocampus, with destinations for either CA1 (from bregma and brain surface, most medial shank: Anterior/Posterior −4.0mm, Medial/Lateral 1.8mm, Dorsal/Ventral −0.5mm) or CA3 (from bregma and brain surface, most medial shank: Anterior/Posterior −3.2mm, Medial/Lateral 2.2mm, Dorsal/Ventral −1.0mm). Group 2 (4 young, 3 aged) received L3 series electrodes above the hippocampus (from bregma and brain surface: Anterior/Posterior −3.2, Medial/Lateral 2.2mm, Dorsal/Ventral −3.5mm) and the lateral entorhinal cortex (not analyzed or discussed in this paper). Both probes were purchased from Cambridge Neurotech. The F-series probes had 64 channels across six shanks, with each shank spanning approximately 150μm and each shank spaced 200 μm from eachother **(Fig. 1A)**. The L3 series probes had 64 channels linearly spaced 50 μm covering 3.15mm and was able to record simultaneously from CA1, CA3, and the dentate gyrus **(Fig. 1B)**.

**Figure 1:**
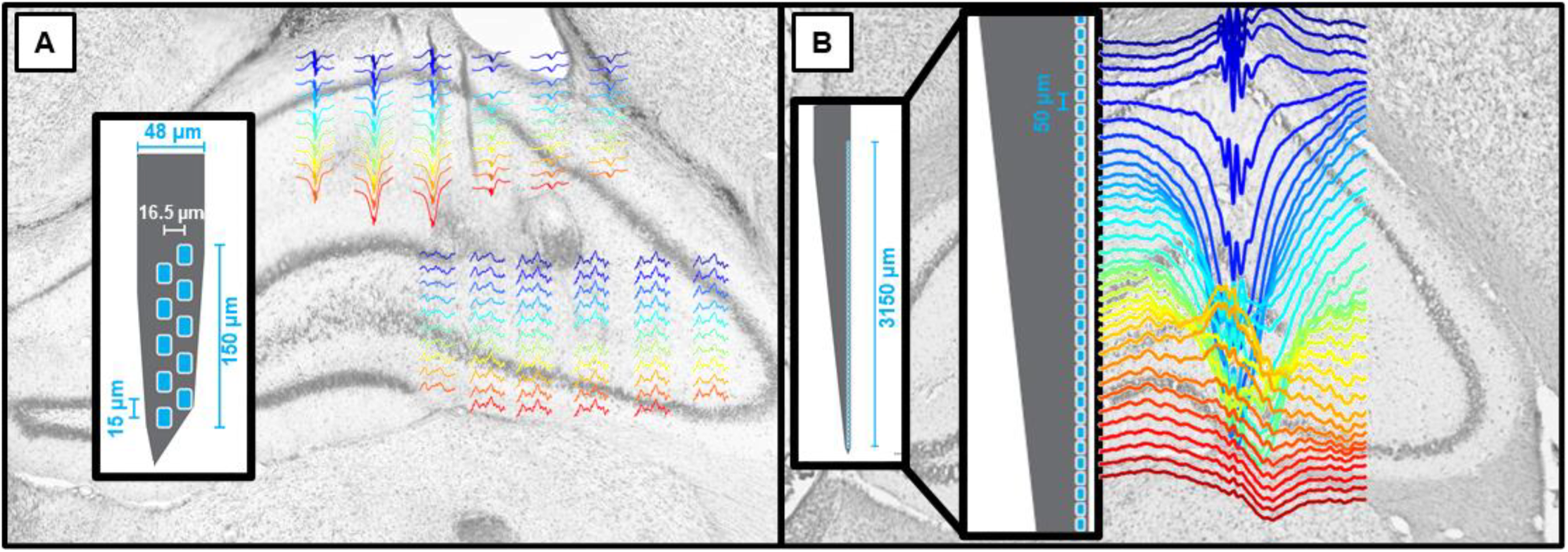
Representative sharp wave/ripples overlayed on hippocampal histology along-side probe array configuration. The change in the direction of the sharp waves, and the power of the ripple, across the hippocampal layers should be noted for both probe types. Rats with two different electrode arrays were included in the current study: one group with dual silicon probes in the CA1 and CA3 sublayers (left), and the other group had a single linear probe spanning 3 mm (right). Color corresponds to the depth of the probes in the brain for a given array, with warmer colors being more ventral.

After a craniotomy was performed at these locations the dura was removed before implanting the probes. A reference electrode was implanted using a steel screw into the cerebellum to remove movement noise. A grounding electrode was placed in the ipsilateral pre-frontal cortex. After implanting, dura-gel was added to the opening to form a flexible, impermeable layer between the air and brain tissue. In-depth surgical instructions can also be viewed on the website for Cambridge Neurotech (https://www.cambridgeneurotech.com/neural-probes). A copper mesh that surrounded the implant to provide both electrical and physical protection was then placed and affixed with dental cement. The ground wire was soldering to the copper mesh cage. Probes were connected to ZIF clips that were mounted in the dental cement and could be attached to clips that ran to a Tucker Davis Technologies electrophysiology recording hardware sampling data at 24,414.0625 samples/second. After surgery, the rats were given 5 mL of warmed, sterile saline for hydration. Simultaneous video recordings were taken from two overhead cameras, one to track LEDs mounted on the ZIF clips to track the animal’s location and head direction, and one infrared camera for post-run behavioral scoring. Raw data was saved to an external server and a copy was saved in a 4 kHz sampling rate format to an additional external hard drive.

### 2.03 Behavioral Procedures for Neurophysiological Recordings

Post-surgery, all animals were re-trained to push the square block prior to performing their respective task (LEGO© discrimination or OPPA training). All rats performed two additional behavioral tasks, an 8-maze alternation and a circular track running task. The 8-maze alternation required the animal to alternate turning direction to receive a reward. Animals completed 50 laps or ran for 60 minutes, whichever came first. The circle task required rats to run in a counterclockwise direction for 50 laps or 60 minutes, whichever came first. All electrophysiology data presented here were analyzed from days in which animals completed the continuous alternation on the 8-maze or circle running. For all recording days, data were collected during 20-minute rest epochs that flanked the behavioral testing. All analyses were restricted to the pre- and post-run recording epochs to assess local field potential dynamics during these ‘offline’ periods.

### 2.04 Detection of High-Frequency Events

To detect sharp wave ripple events, channels were picked that either contained spiking events with burst-firing characteristics, indicating their location in the pyramidal layer, or were chosen based on the direction of the sharp waves. This was easiest to see in the 3.15mm long, L3 probes (**Fig. 2)**, so the power spectral density of all the channels in the F-series probes were compared to those in the L3. Channel selection was the same across both rest epochs but could change across days. Data was then bandpass filtered between 115 and 250 Hz, rectified, and the mean and standard deviation of the amplitude was calculated. Frequency ranges were selected based on a previous investigation of ripples in young and aged rats (Cowen et al., 2020), as well as the ripple frequency distributions in the current dataset **(Fig. 3A-D)**.

**Figure 2:**
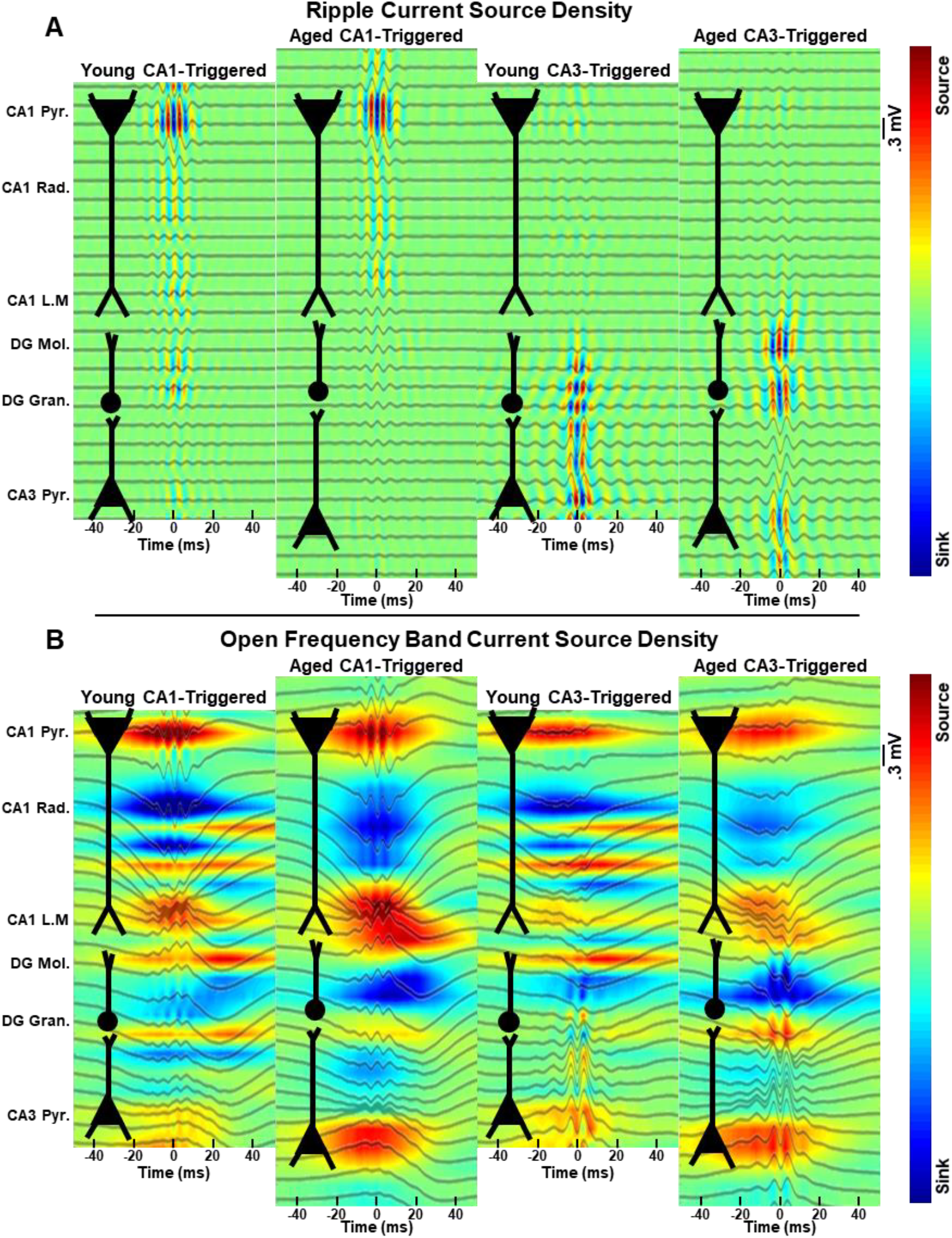
Representative traces and current source densities (CSD) of either ripple filtered **(A)** or unfiltered **(B)** LFP for young and aged rats triggered on ripples in either CA1 or CA3. Traces are shifted to align the CA1 and CA3 layers between rats to show the anatomy aligned with the neurophysiology. Neuronal sub-layers are indicated to the left of the figures. Note the presence of sources and sinks in the pyramidal layer during periods of ripples triggered in the other region (e.g. CA1 ripples during CA3 triggered events). Also evident is the presence of sources and sinks in the molecular layer during pyramidal layer triggered events. Pyramidal layers were identified by spiking activity, radiatum by the first large sink below the CA1 pyramidal layer, lacunosum moleculare and dentate gyrus molecular layer by CSDs triggered to theta (6-12Hz) or gamma (40-100Hz) (respectively) during the run epoch (not shown).

**Figure 3:**
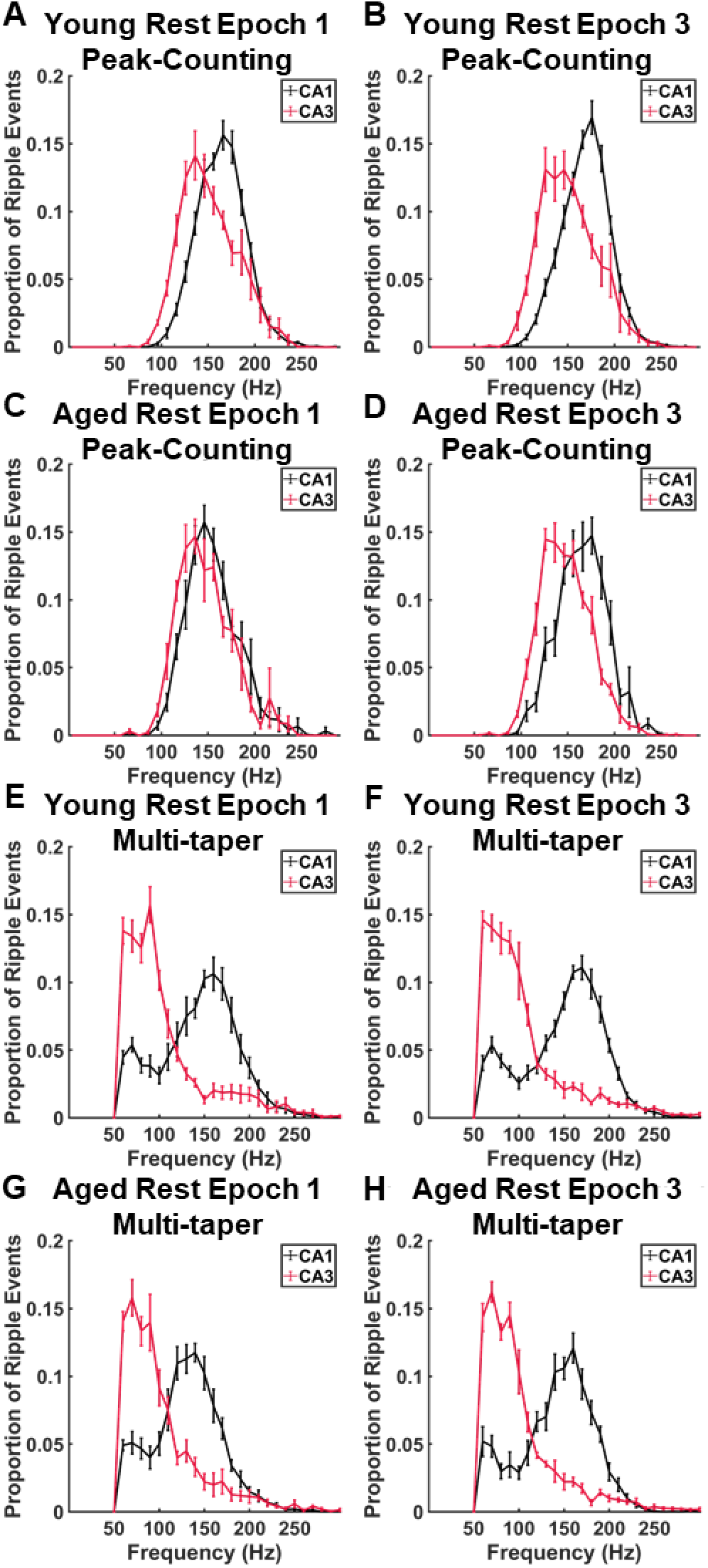
Ripple frequency distributions calculated using peak counting and multi-taper. Ripple frequency distributions were calculated separately for each rat and averaged (young n=8; aged n=7). Error bars are standard error of the means for 10Hz bins. Hartigan’s dip test for bimodality was calculated using 500 bootstraps for each rat. All rats failed to reject the null hypothesis of a unimodal distribution (p>0.05). Additionally, frequency distributions were tested for normality using a one-sampled Kolmogorov-Smirnov test. The null hypothesis of normality was rejected for all rats (p<0.05). Due to the lack of a normal distribution all metrics were calculated using the median for each rat rather than the mean. While other labs have reported a high gamma frequency (90-140Hz) in the CA1 pyramidal layer (Sullivan et al., 2011) this was not found using the peak-counting method used to calculate ripple frequency **(A-D)**. To rule out the possibility of a false negative, ripples were detected, curated, and frequencies calculated using the methods described in Sullivan et al., 2011 **(E-H)**. In short, ripples were detected and curated based on the presence of a spectral peak between 120-200Hz that was significantly larger than the background power spectrum, and frequencies were calculated using a multi-taper method on 100ms windows. Both methods revealed peak frequency differences in CA1 that were higher relative to CA3.

Hartigan’s Test for Bimodality and one-sampled Kolmogorov-Smirnov (KS) tests for distribution normality were computed for each rat’s ripple frequency distribution. Because all rats returned significant KS tests (p<.05) median values for each rat were used for all of the following analyses. Additionally, because all rats returned non-significant Hartigan’s tests for bimodality (p>0.05) no attempt was made to separate ripples based on different frequency ranges. To rule out the possibility that a disproportionate false classification of lower frequency events as ripples in aged compared to young rats, we replicated a previous analysis showing that high frequency hippocampal oscillations were bimodally distributed with a local minima at ~140 Hz, which presumably separated gamma from ripples in CA1 (Sullivan et al., 2011). In short, time series data were filtered between 50 and 250 Hz, and high frequency events with a spectral peak greater than 2 standard deviations above baseline in the 120-200 Hz frequency range were selected for analysis. Frequency was then calculated using multi-taper on 100ms windows. For CA1, the frequency distributions for both young and aged rats had a local minima at 100 Hz, indicating that there was not a disproportionate contamination of gamma in the ripple detections between age groups **(Fig. 3E-H)**.

Ripple detection threshold for the rest epoch ripples was based on thresholds set from the run epoch, as previously described (Maurer et al., 2006). Briefly, a ripple event was detected when the rectified signal crossed 3 standard deviations above the mean. The point where the signal rose and fell above and below 1 standard deviation was then designated the start and end of the ripple event. Ripples events with an interval less than 15ms were combined into one ripple. Ripples shorter than 30 ms or longer than 500 ms were not included in the analysis. To account for noise events such as chewing artifacts the standard deviation of the ripple (115-250 Hz) and sharp wave (14-20 Hz) power across all channels was calculated. If the ripple and sharp wave power standard deviations did not exceed 3 or 5 from the mean, respectively, that ripple was considered a false positive and excluded from further analysis. This approach capitalizes on the fact that the localized current sources and sinks within the laminar structure of the hippocampus produce changes in the measured LFP signal across different laminar positions, while noise is typically homogenous across all channels. Ripple coincidence between CA1 and CA3 was measured from the peak power of the ripple due to the start of a ripple being highly variable based on what thresholds are used.

### 2.05 Ripple Quantification

All ripple quantification was done on the median values of the respective metric. Sample sizes were based on rat count, not ripple numbers, as to avoid violating the assumption of independent samples and inflating statistical power. Seven ripple metrics were calculated: frequency, power, amplitude, length, and envelope half-width time length. Amplitude and envelope half-width time length are reported in **Supplemental Figure 1**, as they were consistent with power and length measures, respectively. Frequency was calculated based on the number of local minimums divided by the length of the ripple. Ripple power was calculated as the sum of the squared values of the filtered (115-250 Hz) signal divided by the ripple length. Ripple length was described earlier as the time difference between when the envelope of the filtered, rectified signal fell below 1 standard deviation of the mean. Ripple amplitude was the maximum absolute value of negative values of the filtered (115-250 Hz) signal. Ripple envelope half-amplitude time width was the time length of the negative envelope at half its maximum amplitude.

### 2.06 Sharp Wave Quantification

Sharp waves were only quantified on the rats in Group 2 that received the L-series electrode arrays (4 young, 3 aged) because they were implanted with the 3.15mm long probe, which spanned all hippocampal sublayers. The radiatum channel was chosen using the current source density (CSD) during CA1 ripple events that co-occurred with CA3 ripple events. The channel with the largest sink located above the dentate gyrus was considered as the radiatum. This was hand verified using the raw traces. Sharp waves were not detected independently of ripple events. Power was calculated on the sum of the squared raw trace of that channel during ripple events. Power was calculated on the raw traces because the transient nature of sharp waves meant that filtering the signal to a narrow band caused the formation of artificial oscillations on either side of the sharp-wave event.

### 2.07 Principal Component Analysis and Correlation Testing

Principal Component Analysis (PCA) was performed in SPSS on the three different ripple metrics of frequency, power, and length. Values for each rat were derived from the median of the combined day 1 and day 2 ripples. Epochs were kept as separate data points. This was due to the consistent significance due to rest epoch, but lack of significant differences across days. There were therefore two values for each rat/metric combination. Significant loadings in the PCA matrix were considered those greater than 0.4. Metrics that loaded significantly onto the same component were then tested for correlations based on their Pearson’s Correlation Coefficient and t-statistic using the MATLAB function *corrcoef*. Young and aged rats were analyzed separately as well as combined. Correlations on the raw ripple data, where every ripple event was its own data point, were also calculated. This was done for verification purposes only and significance testing was not performed on this dataset due to the overpowered nature of the analysis and violation of independence of samples.

### 2.08 Ripple Rate and Rat Speed

Ripple rates (ripples/second) were calculated as the number of ripple occurrences in a rest epoch over the length of the rest epoch. Average rat speed (cm/second) was also calculated to ensure that potential differences in ripple occurrences were not due to rat activity differences in the resting pot. For illustrative purposes, ripple rate and rat speed were also binned into two-minute blocks **(Supplemental Figure 2)**.

### 2.09 Statistical Analysis

Mixed Model Analysis of Variances (ANOVA) were used to analyze age differences across multiple rest periods, days, and condition for ripple quantification metrics with two between subject factors of age group (young and aged) and familiarity (novel day 1 and familiar day 2). Rest epoch (pre-run rest epoch 1 and post-run rest epoch 3) was the within subjects factor. The CA1 and CA3 regions were analyzed separately. A rejection of the null hypothesis was considered with an alpha level of p<0.05. This design was also used to study ripple co-occurrence and rat speed. To study ripple metrics as a function of the co-occurrence between CA1 and CA3 ripples, the statistical design was again a mixed model ANOVA but with two between subjects factors of age group and rest epoch, and one within subjects factor of ripple co-occurrence (non-co-occurrence and with co-occurrence). Days and regions were analyzed separately, and significance was set at p<0.05. Ripple rates were tested for significance using a mixed model design with a between subjects factor of age and region, and a within subjects factor of rest epoch. Days were analyzed separately. Finally, ripple rates and rat speeds (binned into 2-minute time windows) were examined with a Mixed Model ANOVA with between subjects factor of age, and the within subjects factor of time blocks (df=9). Rest epochs and days were all analyzed separately.

### 2.10 Histological Probe Verification

After all experiments were completed, rats were injected with 0.9mL of SomnaSol (pentobarbital) and perfused through their left ventricle with 4% para-formaldehyde (PFA). Brains were extracted and suspended in more PFA until cryoprotection with 30% sucrose PBS solution. Once brains lost buoyancy, they were flash frozen with dry ice and embedded in Tissue-Tek Optimal Cutting Temperature compound. Brains were then stored in a −80°C freezer overnight for temperatures to stabilize. The next day the brains were moved to a −20°C freezer overnight. Brains were sectioned onto Fisher Superfrost Plus slides in 50μm thick slices in a Cryostat kept at −20°C. These slides were stained with Nissl and imaged at 20x magnification for verification of probe placement.

## 3 RESULTS

### 3.1 CA1, but not CA3, ripple metrics changed significantly between rest epochs and age groups

Ripple frequency, power, and length were not normally distributed. Thus, statistics were calculated based on the median of ripple events found within a rest epoch. In both age groups, CA1 ripple frequency, power, and length significantly increased from pre- to post-run rest epochs **(Fig 4C, E, G;** F_[1,26]_>14.73, p<0.001, Mixed Model ANOVA; see **Table 1** for summary of all statistics). This increase in post-run rest ripple metrics was significantly larger in the aged rats as indicated by a significant interaction of age x rest epoch for ripple frequency (F_[1,26]_=4.42, p<0.05) and length (F_[1,26]_=7.75, p<0.01) **(Fig. 4C, G)**. Consistent with a previous report (Wiegand et al., 2016), aged CA1 ripples had significantly reduced frequency compared to their younger counterparts (F_[1,26]_=4.90, p<0.05). To ensure rigor, ripple amplitude and ripple envelop half-amplitude time were also calculated as proxies for power and length, respectively. Consistent with the previous analyses, ripple amplitude was significantly larger during the post- compared to pre-run rest epoch (F_[1,26]_=16.46, p<.001), and ripple envelope half-amplitude time had a significant interaction of rest epoch and age (F_[1,26]_=5.02, p<.05). These metrics can be seen in **Supplemental Figure 1**. CA3 ripple events were found independently of CA1, but frequency thresholds (115-250 Hz) were kept the same. No significant differences between groups or rest epochs were detected (F_[1, 26]_<4.22, p>.05 for all comparisons) **(Fig. 4D, F, H) (Table 1)**.

**Figure 4:**
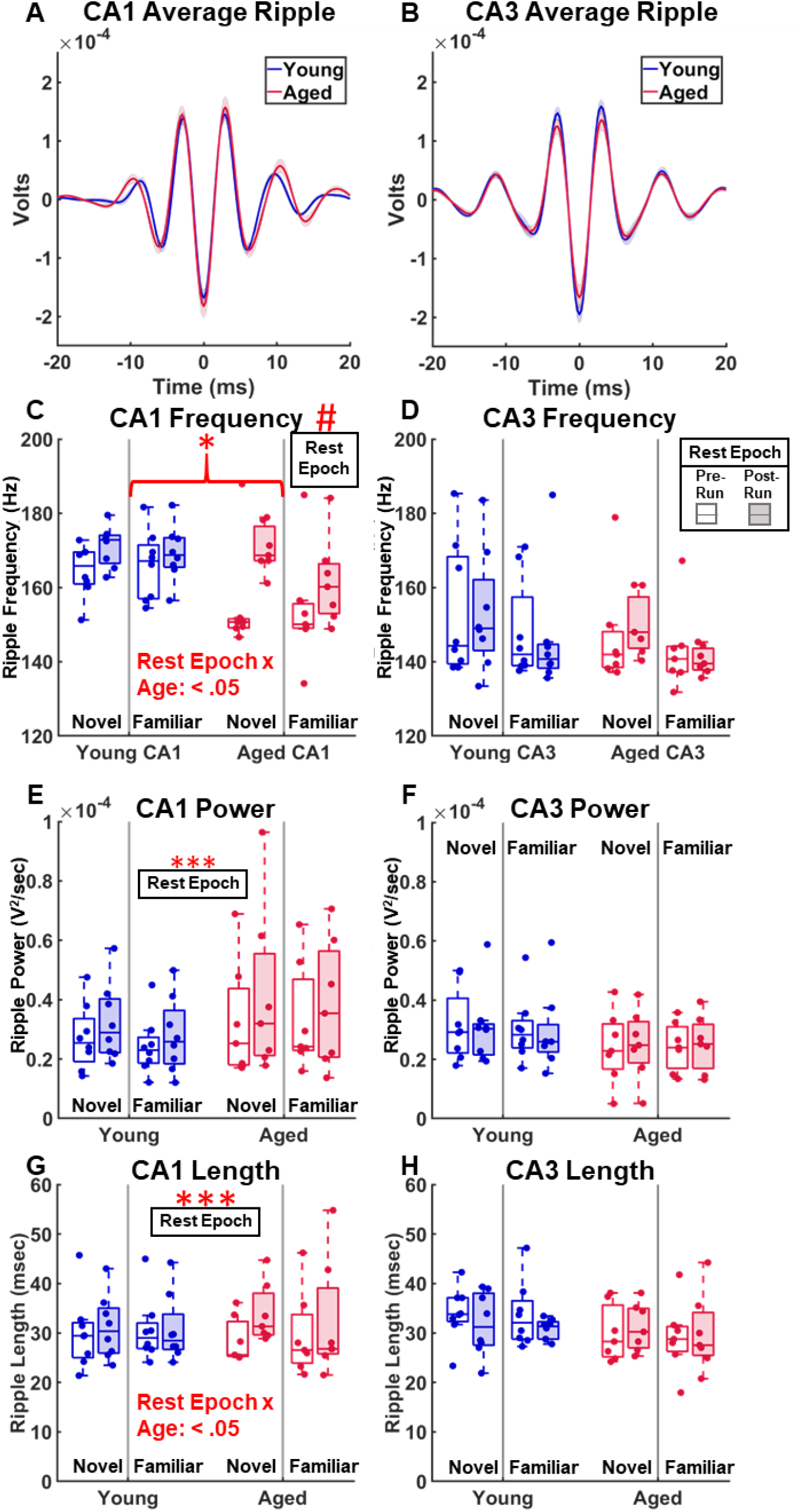
Young and Aged CA1 and CA3 Ripple Quantification. Ripple quantification in the CA1 and CA3 subregions of the hippocampus of young and aged rats. **(A-B)** Averaged, overlayed, filtered ripples. The peaks of the young and aged CA1 average ripples diverge while those in CA3 do not. **(C, D)** Ripple frequency separated by region, age, day, and rest epoch. CA1 ripples were significantly different between age groups (F_[1,26]_=4.90, p<.05), and rest epoch (F_[1,26]_=27.8, p<.0001). The rest epoch and age interaction was also significant (F_[1,26]_=4.42, p<0.05). CA3 ripple frequency had no significant differences between age groups, epoch or day. **(E-F)** CA1 ripple power had significant variation between different rest epochs (F_[1,26]_=15.85, p<.001). No effect was seen in CA3 ripple power. **(G-H)** CA1 Ripple length varied significantly by rest epoch (F_[1,26]_=14.73, p<.001), and there was a rest epoch and age interaction (F_[1,26]_=7.76, p<.01). No effects were seen in CA3.

**Table 1:** Statistical Results of Ripple Quantification Metrics Shown in Figure 4.

| Factor | CA1 |  |  | CA3 |  |  |
| --- | --- | --- | --- | --- | --- | --- |
|  | Frequency | Power | Length | Frequency | Power | Length |
| Age | $F_{[1,26]}=4.90$<br>$p=.04$ * | $F_{[1,26]}=1.81$<br>$p=.19$ | $F_{[1,26]}=.028$<br>$p=.87$ | $F_{[1,26]}=1.36$<br>$p=.25$ | $F_{[1,26]}=1.92$<br>$p=.18$ | $F_{[1,26]}=1.57$<br>$p=.22$ |
| Day | $F_{[1,26]}=0.61$<br>$p=.44$ | $F_{[1,26]}=.14$<br>$p=.71$ | $F_{[1,26]}=.0008$<br>$p=.98$ | $F_{[1,26]}=2.01$<br>$p=.17$ | $F_{[1,26]}=.01$<br>$p=.90$ | $F_{[1,26]}=.18$<br>$p=.68$ |
| Age x Day | $F_{[1,26]}=0.50$<br>$p=.49$ | $F_{[1,26]}=.04$<br>$p=.84$ | $F_{[1,26]}=.0046$<br>$p=.95$ | $F_{[1,26]}=.0003$<br>$p=.99$ | $F_{[1,26]}=.035$<br>$p=.85$ | $F_{[1,26]}=.003$<br>$p=.96$ |
| Rest Epoch | $F_{[1,26]}=27.80$<br>$p=1.6e-5$ # | $F_{[1,26]}=15.85$<br>$p=.0005$ *** | $F_{[1,26]}=14.73$<br>$p=.0007$ *** | $F_{[1,26]}=.09$<br>$p=.77$ | $F_{[1,26]}=.005$<br>$p=.94$ | $F_{[1,26]}=.77$<br>$p=.39$ |
| Age x Epoch | $F_{[1,26]}=4.42$<br>$p=.045$ * | $F_{[1,26]}=.99$<br>$p=.33$ | $F_{[1,26]}=7.76$<br>$p=.0098$ ** | $F_{[1,26]}=.06$<br>$p=.81$ | $F_{[1,26]}=.99$<br>$p=.33$ | $F_{[1,26]}=4.22$<br>$p=.0501$ |
| Day x Epoch | $F_{[1,26]}=2.90$<br>$p=.10$ | $F_{[1,26]}=.99$<br>$p=.33$ | $F_{[1,26]}=2.24$<br>$p=.15$ | $F_{[1,26]}=.63$<br>$p=.43$ | $F_{[1,26]}=2.52e-5$<br>$p=1.00$ | $F_{[1,26]}=.11$<br>$p=.75$ |
| Age x Day x Epoch | $F_{[1,26]}=.45$<br>$p=.51$ | $F_{[1,26]}=.21$<br>$p=.65$ | $F_{[1,26]}=.52$<br>$p=.48$ | $F_{[1,26]}=.16$<br>$p=.69$ | $F_{[1,26]}=.002$<br>$p=.96$ | $F_{[1,26]}=.01$<br>$p=.92$ |
| *: $p<.05$ **: $p<.01$ ***: $p<.001$ #: $p<.0001$ | | | | | | |

### 3.2 CA1 and CA3 ripple metric relationships

To determine the patterns of association among the ripple metrics of frequency, power, and length in CA1 and CA3, ripple metrics during the pre-run and post-run epochs for both subregions were subject to principal component analysis (PCA) **(Table 2)**. A PCA with a varimax rotation was performed due to the occurrence of split loadings without the rotation. The rotation converged in iterations. This analysis revealed no problematic redundancies for any task variables (problematic redundancy was considered a loading of >±0.2 across multiple components). According to total variance explained, a model using the top two components (eigenvalues >1.0) explained 59.36% of the variance in the ripple metrics, with the first component accounting for 41.45% of the variance and the second component accounting for 17.91% of the variance. Item communalities were moderate to high (ranging from 0.20 to 0.85 for all items). CA1 length and CA3 length both loaded positively onto the first component (0.83 and 0.85, respectively), suggesting that ripple length across hippocampal subregions was related. CA1 power, CA3 frequency, and CA3 power negatively loaded onto the first component (−0.60, −0.51, and −0.49, respectively). These three variables positively loaded onto the second component (0.64, 0.20 and 0.32, respectively), however. Interestingly, CA1 frequency showed a high negative loading (−0.86) onto the second component.

**Table 2:**
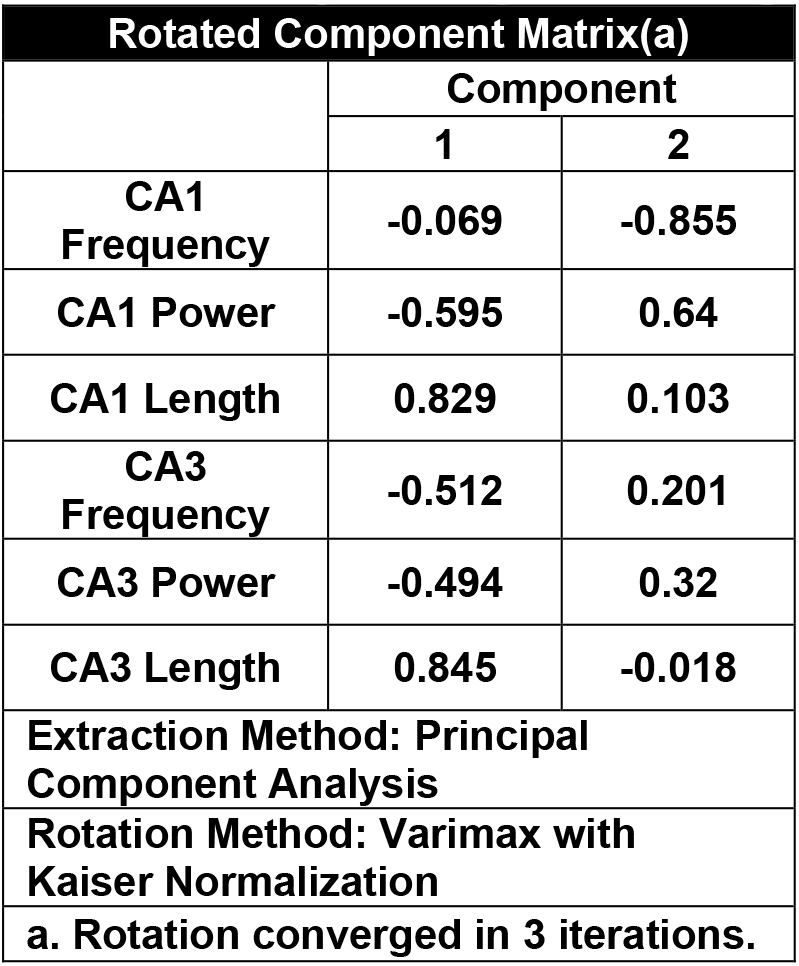
Principal Component Analysis for CA1 and CA3 ripple metrics.

Correlation values of ripple metrics that loaded onto the same components were then determined with a Pearson’s correlation coefficient and its accompanying t-statistic for any regions and metrics that loaded greater than 0.4 onto their respective PCA components. Due to the consistent, significant effect of rest epoch across CA1 ripple metrics, but not of training day/familiarity, rest epoch values were analyzed separately, and days were combined (i.e., one data point would be the median value of all ripple events that occurred for the pre-run rest period on both days of training). Metrics that loaded in the same direction (positive or negative) on the PCA component tended to correlate positively with each other. CA1 power and CA3 frequency were significantly correlated for the young rats (r_[14]_=.611, p<.05), aged rats (r_[12]_=.623, p<.05), and when the young and aged groups were combined (r_[28]_=.474, p<.01) **(Fig. 5A)**. In addition to plotting the median ripple metric values, the raw data for each independent ripple event are displayed as an inset for each correlation to illustrate that the correlation trend was similar between the raw and summary data. Statistics were not calculated for these due to having too high of statistical power from the large sample sizes and a nested design. Interestingly, only the aged CA1 and CA3 ripple power were correlated (r_[12]_=.758, p<.01) **(Fig. 5B)**. The ripple power in the young group in CA3 appeared to have a small variance across rats and rest epochs, which may have contributed to insufficient parametric space for detecting a significant power correlation between CA1 and CA3. The positive correlation was also qualitatively evident in the raw, inset figure. Because ripple power in both CA1 and CA3 did not significantly vary between age groups, it is likely that the inability to detect a significant CA1-CA3 correlation in ripple power of young rats was due to the small sample size.

**Figure 5:**
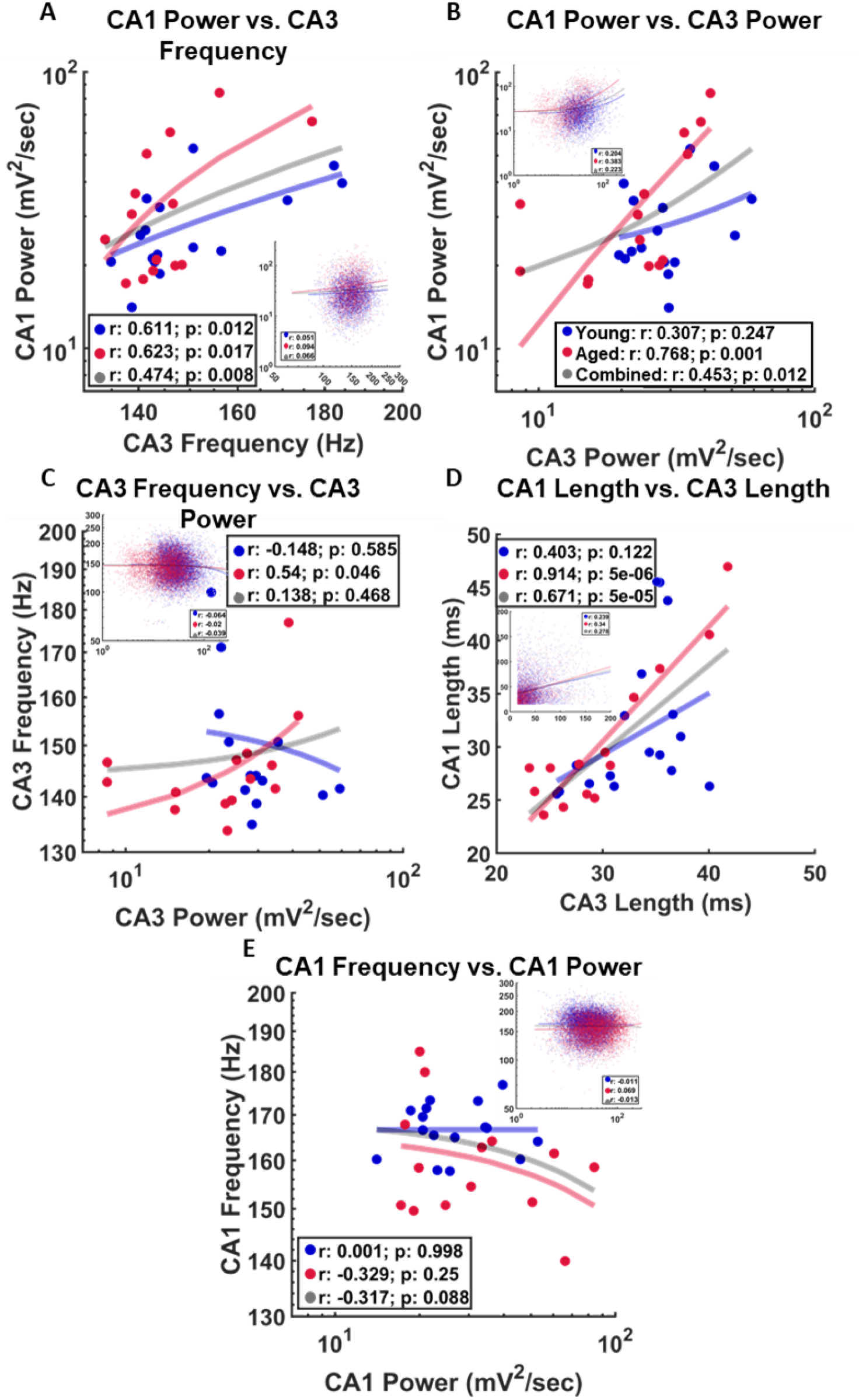
Ripple Metric Relationships. **(A)** CA1 power and CA3 frequency were significantly correlated for both young (r_[14]_=.611, p<.05) and aged rats (r_[12]_=.623, p<.05), as well as the two groups combined (r_[28]_=.474, p<.01). **(B)** CA1 power and CA3 frequency also significantly correlated, but only for aged rats (r_[12]_=.768, p<.001), and when young and aged rats are combined (r_[28]_=.453, p<.05). **(C)** CA3 power and CA3 frequency were significantly correlated for aged rats (r_[12]_=.54, p<.05), but not the young (r_[14]_=−0.148). **(D)** There was a significant correlation between CA1 and CA3 ripple length in aged rats (r_[12]_=.914, p=5e-6) and when both age groups were combined (r_[28]_=.671, p=.5e-5). For the young group, however, while a positive relationship was evident in the raw and summary data, this did not reach statistical significance (r_[14]_=0.403, p>.05). **(E)** CA1 power and CA1 frequency were the only two region/metric combinations that loaded onto component 2 of the PCA (>0.4). They were inversely loaded, but there was no significant negative correlation between the metrics for either young, aged, or combined groups (p>0.5). Power and Frequency are plotted on a log scale.

CA1 and CA3 ripple length strongly loaded onto component 1 (0.829 and 0.845, respectively). This was likely carried by the significant correlation between CA1 and CA3 ripple length in the aged rats (r_[12]_=0.914, p=5e-6). This relationship did not reach statistical significance in the young rats (r_[14]_=0.403, p=0.122) **(Fig. 5D)**. The correlation between lengths was qualitatively evident in the raw data for both age groups. Thus, it is possible that the lack of a significant correlation in the young rats between CA1 and CA3 ripple length was related to the small sample size. CA1 frequency loaded negatively and CA1 power loaded positively onto component 2 of the PCA. A relationship between these variables, however, was not significant for either young (r_[14]_=0.001, p=0.998) or aged (r_[12]_=−0.329, p=0.25) rats, or when the two groups were combined (r_[28]_=−0.317, p=0.088).

### 3.3 Behavior differentially impacts the co-occurrence of CA1 and CA3 ripples in young and aged rats

Due to the decreased synaptic efficacy of the Schaffer collaterals (Landfield et al., 1986; Barnes et al., 1992; Deupree et al., 1993; Tombaugh et al., 2002), it is conceivable that aged rats would exhibit a decreased ability of CA3 to drive coordinated activity in CA1. To test for this, ripple co-occurrence was examined by measuring the cross-region inter-ripple interval with respect to CA1 ripples (**Fig. 6**), and with respect to CA3 ripples (data not shown). The timing for each ripple was based on the peak amplitude of the filtered and rectified signal. A CA1-CA3 ripple co-occurrence was considered to take place when the ripple peaks fell within a ±50 ms window, and the summed value between these time points was used for statistical testing. In the aged rats, CA1-CA3 ripple co-occurrence was lower prior to running on the track, but increased after running. This pattern was not observed in the young rats, which was indicated by a significant interaction between rest epoch and age for ripple co-occurrence measured relative to CA1 ripples (F_[1, 26]_=6.08, p=.02). Importantly, similar results were obtained when CA1-CA3 ripple occurrence was measured with respect to CA3 ripples (F_[1,26]_=4.27, p=.049).

**Figure 6:**
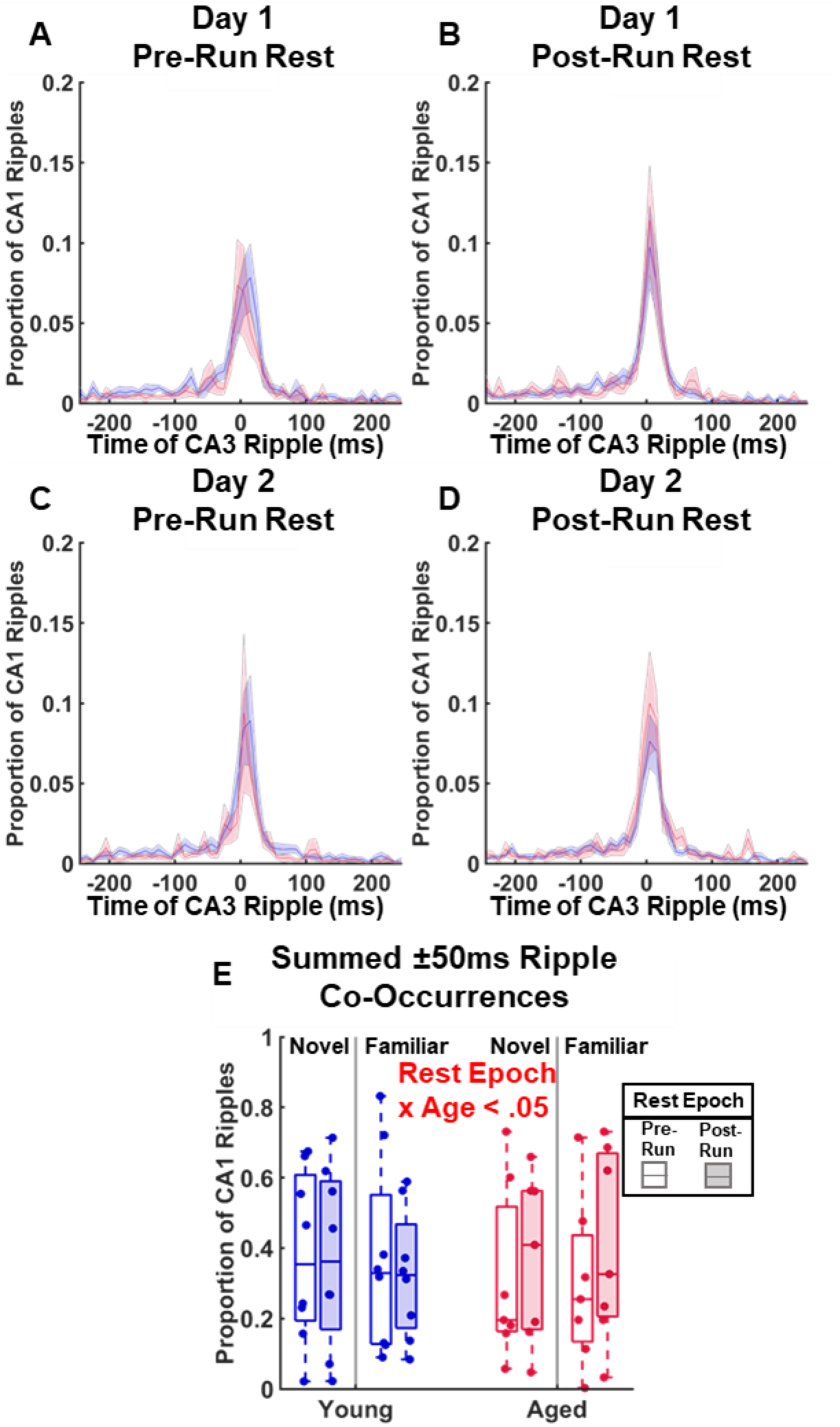
CA1 Ripple Co-Occurrence with CA3 by Day, Rest Epoch, and Age. **(A-D)** Average histograms (with standard error of the means) of the proportions of CA1 ripples that had a corresponding CA3 ripple. Ripple timing was measured from the peak power of the respective ripple. Global maximums tended to occur between ±50 ms. **(E)** There was a significant interaction of rest epoch and age (F_[1, 26]_=6.08, p=.02).

Based on the finding of a significant interaction effect of age and rest epoch on CA1-CA3 ripple co-occurrence, it was hypothesized that changes in age-related synaptic efficacy could manifest as differences between the characteristics of ripples that coincided with a ripple from the other region. Quantified ripple metrics were therefore separated into co-occurring **(Co)** and non-co-occurring **(Non)**. Except for CA1 frequency on day 2 **(Fig. 7Aii)**, all ripple metrics for both CA1 and CA3 (frequency, power, and length) were significantly greater when they co-occurred during both days of testing (F_[1,26]_>9.43, p<0.01, see **Table 3** for all comparisons) **(Fig. 7)**. This effect only significantly interacted with age for CA1 ripple length on day 2 (F_[1,26]_=4.76, p=.04) **(Fig. 7Cii)**. Age had a significant effect on CA1 ripple frequency for both day 1 (F_[1,26]_=6.62, p=0.017) and day 2 (F_[1,26]_=5.42, p=.028) **(Fig. 7A)**. On day 1, there was also a significant effect of rest epoch (F_[1,26]_=27.64, p=1.71e-5) and an interaction of rest epoch and age (F_[1,26]_=6.47, p=0.017). While co-occurrence significantly resulted in ripples with higher power (**Fig. 7C)** and longer duration (**Fig. 7E)**, this increase did not significantly interact with rest epoch for Day 1 or Day 2 (F_[1,26]_<0.60, p>0.45 for all comparisons). Co-occurrence effects on CA1 ripple power and length also did not significantly interact with age, with the exception that on Day 2 the increase in CA1 ripple length between co-occurring and non-co-occurring ripples was significantly smaller for the aged compared to young rats (F_[1,26]_=4.76, p<0.05). When the same analysis was run for CA3 ripples, there was again a significant increase in CA3 ripple frequency, power, and length when the ripple coincided with a CA1 ripple (**Table 4**; p<0.01 for all comparisons). Age and rest, however, did not show a significant interaction with co-occurrence for any CA3 ripple metric (p>.05 for all comparisons) (**Fig. 7B,D,F**).

**Figure 7:**
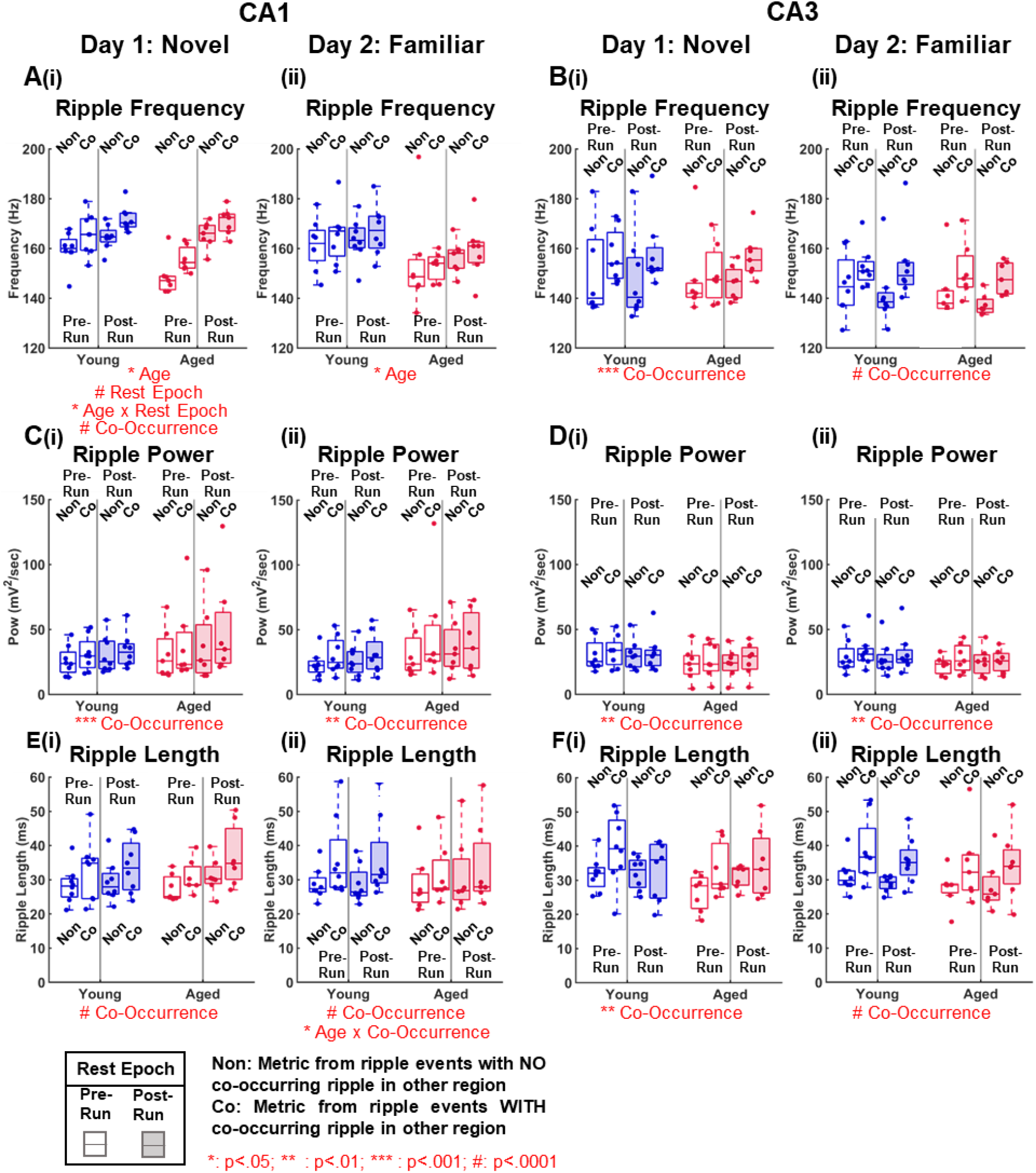
Ripples that co-occur with a ripple in the other region have increased frequency, power and length across age groups. CA1 and CA3 ripple metrics reported in Fig. 4 were examined based on whether or not a ripple co-occurred with a ripple in the other region. Ripples that co-occurred with other ripples are labeled **(Co)**, and those that do not are labeled **(Non)**. Ripples were also separated by day and pre/post run rest epoch. Except for CA1 frequency on day 2 **(Aii)**, all metrics were significantly affected by co-occurrence versus not (F_[1,26]_>9.43, p<0.01). CA1 ripple frequency on both days 1 and 2 **(A)** had a significant interaction with age (F_[1,26]_>5.42, p<0.05). CA1 ripple frequency on day 1 was significantly affected by rest epoch (F_[1,26]_=27.64, p<0.0001), and had a significant age x rest epoch interaction (F [1,26]=6.47, p<0.05). **(Ei)** An interaction between co-occurrence and age was also significant on day 2 (F_[1,26]_=4.76, p<0.05). There was no effect of either rest epoch, age, or interaction (F_[1,26]_<2.89, p>0.05) for CA3 ripples **(B,D,F)** on either day 1 or day 2 for any metric.

**Table 3:**
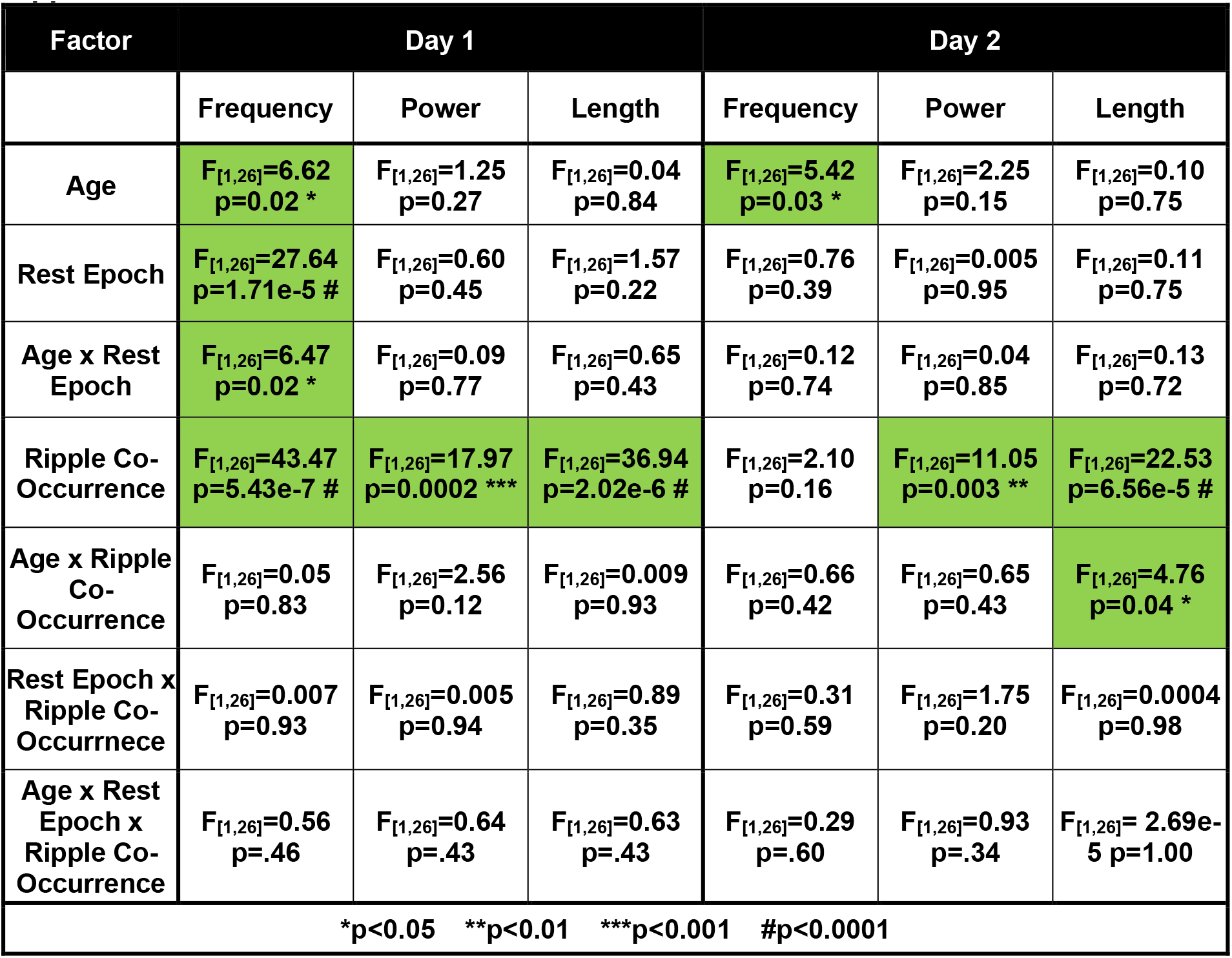
Statistical Results for CA1 Ripple Metrics Separated by Co-Occurrence with CA3 Ripples.

### 3.4 Sharp wave power was significantly impacted by rest epoch and co-occurrence, but not age

Due to the widely reported decrease in synaptic efficacy at the CA3 to CA1 Schaffer collateral synapse, sharp wave power in the stratum radiatum was examined for a potential age difference. Sharp waves could only be quantified in rats that were implanted with a long electrode array. Thus, only rats in Group 2 with the long linear silicon probe (Young n=4; Aged n=3) were included. The radiatum was determined based on the channel with the peak power in the current source density analysis of the wide-band signal during ripple events **(Fig 8A)**. Interestingly, this was often located in the dentate gyrus layer, so all channels were hand verified and corrected if found to be erroneously selected. Radiatum power was calculated based on the summed square of filtered (4-30 Hz) signal from the chosen channel during a CA1 ripple event and the median value for each rat was used for statistical analysis **(Fig. 8B, C)**. Sharp waves that co-occurred with a CA3 ripple were labeled **(Co)**, and those that did not were labeled **(Non)**. There was a significant increase in sharp wave power when the depolarization coincided with a CA3 ripple, compared to when no ripple was detected for Day 1 (F_[1, 10]_=7.45, p<0.05) but not for Day 2 (p>0.05). There was also a significant interaction between co-occurrence and rest epoch on Day 1 sharp wave power (F_[1, 10]_=7.22, p<.05). The factors of co-occurrence, and rest epoch on sharp wave power did not significantly interact with age (p>0.05 for all comparisons).

**Figure 8:**
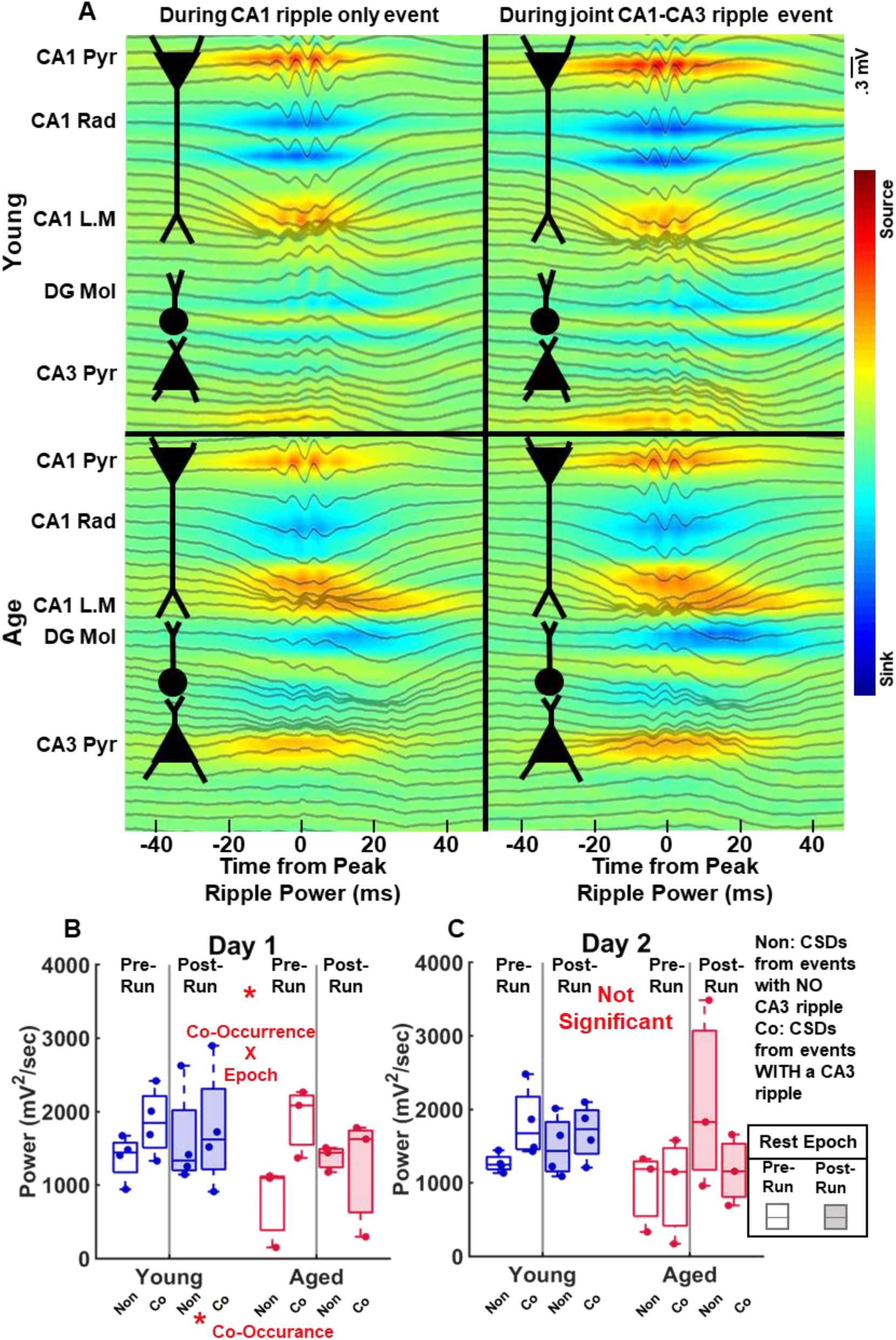
Radiatum Sharp Wave Power separated by co-occurrence. **(A)** Current source densities from a representative young and aged rat, separated by whether the CA1 ripple was detected within ±50 ms of a CA3 ripple. Black lines are the averaged raw LFP traces and a simplified neuron diagram is included for showing how the ionic sources and sinks align with the hippocampal layer anatomy. Sources (red) are heaviest in the radiatum, but interestingly also near the dentate gyrus granular layer. Qualitatively, radiatum power was larger when CA1 ripples co-occurred with a CA3 ripple. **(B-C)** Co-occurrence was only significant for day 1 (F_[1,10]_=7.45, p<0.05). The interaction of co-occurrence and rest epoch was also significant for day 1 (F_[1,10]_=7.22, p<0.05).

**Figure 9:**
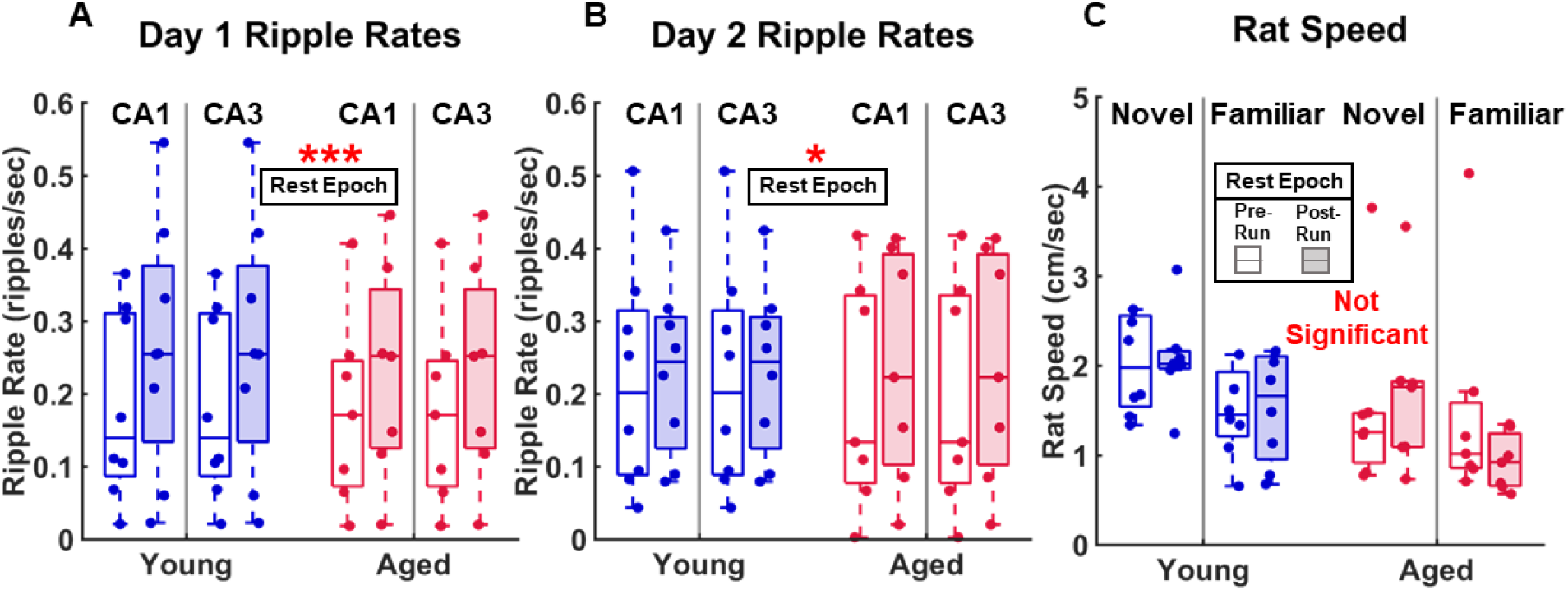
Ripple Rate and Rat Speed. **(A,B)** Median ripple rates for day 1 and day 2 by rest epoch for CA1 and CA3. There was a significant effect of rest epoch for both day 1 (F_[1,26]_=18.12, p=.0002) and day 2 (F_[1,26]_=7.40, p=.012), with the post-run rest epoch having a higher occurrence of ripples for both regions than during the pre-run rest epoch. **(C)** Rat speed calculated for both days and rest epochs. There were no significant effects or interactions (F_[1,26]_<2.07, p>.05).

### 3.5 CA1 and CA3 ripple rates were significantly affected by rest epoch but not age

Ripple rates were calculated by dividing the number of ripples in a rest epoch by the rest epoch time. Ripple rate significantly increased between the pre- and post-run rest epochs in both CA1 and CA3 for both Day 1 (F_[1,26]_=18.12, p=.0002) and Day 2 (F_[1,26]_=7.40, p=.012). Finally, rat speed was analyzed to minimize the possibility that varying activity levels were a contributing factor to the observed ripple changes. No significant effects or interactions were found for age or rest epoch (F_[1,26]_<2.07, p>.05).

## 4 DISCUSSION

The current study documented ripple characteristics and dynamics in both the CA1 and CA3 hippocampal subregions of young and aged rats. The primary findings were that ripple frequency, and length during the pre-run rest epoch were significantly reduced in aged compared to young rats in CA1 only. Interestingly, ripple frequency, power, and length increased between the pre- and post-run rest epochs, and for both frequency and length this increase was more evident in the aged compared to young rats. In contrast to CA1, there were no significant changes in ripple frequency, power, or length in CA3 between age groups or rest epoch. When ripples across subregions were compared, CA3 ripple power and length were related to CA1 power and length, and higher frequency CA3 ripples were also associated with increased ripple power in CA1. These relationships were consistent across age groups. While there was not a significant effect of age on the probability that CA1 and CA3 ripples would co-occur within ±50 msec of each other, there were notable differences in both CA1 and CA3 ripple characteristics when co-occurrence was detected versus when it was not. Specifically, in both age groups frequency, power, and length were all increased in co-occurring ripples compared to when co-occurrence was not detected.

Previous work has reported that in CA1 the frequency of the ripple oscillation is significantly decreased in aged compared to young rats *in vivo* (Wiegand et al., 2016; Cowen et al., 2020), and in slices obtained from young or aged rats (Kouvaros et al., 2015). We replicate this observation and extend it to show that within CA3 there were no significant differences in ripple frequency, power, or length between young and aged rats. The latter finding is somewhat surprising considering the well documented increase in excitability of CA3 pyramidal neurons (Wilson et al., 2005a; Robitsek et al., 2015; Simkin et al., 2015; Thomé et al., 2016; Maurer et al., 2017; Lee et al., 2021) and the disruptions in CA3 inhibition in advanced age (Shetty and Turner, 1998; Shi et al., 2004; Stanley and Shetty, 2004; Spiegel et al., 2013; Thomé et al., 2016). One explanation for this the discordant findings between CA3 single-unit firing and ripples is that CA3 hyperactivity is only something that manifests during waking behaviors and not during periods of rest.

While the current study replicated previous reports of reduced ripple frequency with age in CA1, there were no age differences in CA3 ripples. This observation suggests that age-associated CA1 ripple differences are either due to changes within the small network dynamics of CA1, or related to afferent input alterations in aged animals from other regions that contribute to ripple generation. One possible explanation for age-related reduction in ripple frequency is the reported increase in gap junctions with age (Barnes et al., 1987). In a recent network model, a higher probability of gap junctions was demonstrated to lead to a slight reduction in ripple frequency (Holzbecher and Kempter, 2018). This idea should be directly tested by blocking gap junctions in CA1 and measuring the impact on ripple characteristics.

The overall relatively low probability of CA1 ripples not co-occurring with a CA3 points to other regions being important for CA1 ripple generation. While CA1 ripples are typically considered to be initiated by spontaneous CA3 pyramidal cell firing, other mechanisms of CA1 ripple generation have been reported, such as excitatory input from the subiculum (Imbrosci et al., 2021) or CA2 (Oliva et al., 2016) leading to synchronous depolarization in CA1. To date, the electrophysiological properties of the subiculum and CA2 across the life span are not known and should be examined moving forward. Importantly, there was no main effect of age on CA3-CA1 ripple co-occurrence. Relatedly, sharp wave power in the CA1 *stratum radiatum* was comparable between young and aged rats **(Fig. 8B)**, which is consistent with *in vitro* neurophysiological observations in young and aged mice (Kanak et al., 2013). These data suggest that CA1-CA3 coordination during rest is largely functionally preserved in advanced age. While the probability of CA3-CA1 ripple co-occurrence did not vary with age, there was a significant interaction between age and rest epoch on the proportion of CA1 ripples that co-occurred with a CA3 ripple **(Fig. 6E)**. Specifically, there was an increase in ripple co-occurrence from pre- to post-run rest in the aged group, but not the young. Because it is reported that aged rats have fewer functional synapses (Barnes et al., 1992; Nicholson et al., 2004) and impaired temporal summation at the CA3 to CA1 Schaffer collateral synapse (Rosenzweig et al., 1997), it is conceivable that the behavioral episode was able to promote enhanced CA3-CA1 coordination in the aged rats. Another, not mutually exclusive, possibility is that the change in ripple co-occurrence in aged rats could reflect a shift in subiculum generated ripples (Imbrosci et al., 2021) before behavior to more CA3 generated ripples following behavior.

While CA3 ripples did not vary with age, there were notable differences in the characteristics of CA1 ripples that co-occurred within 50 ms of a CA3 ripple versus those that did not. CA1 ripple power, frequency, and length were all increased when co-occurring with a CA3 ripple, and this was consistent across age groups. This observation suggests that CA1 ripples generated from CA3 are distinct from those generated by the subiculum. Alternatively, it is also possible that CA1 ripples that did not co-occur with a CA3 ripple were ripples that traveled along the septotemporal axis of CA1, rather being locally generated. A previous study reported that ripples that propagate along the septotemporal axis lose frequency and amplitude (Patel et al., 2013).

### 4.1 Limitations

The current study has two primary limitations. First, the experimental design did not directly allow ripples to be related to cognitive performance as rats were not participating in a hippocampus-dependent behavior during neurophysiological recordings. Second, while the 64-channel longitudinal recording shanks enabled current source density analysis for channel localization, they did not permit single unit spiking activity from individual neurons to be recorded. As a result, the ripple analyses could not be complimented by neuron spiking data. Thus, it is not possible to offer further explanations as to why CA3 hyperactivity in aged animals is not reflected in the rest-related ripple activity. Future investigations will be necessary to tease apart the relationship between CA3 neuron spiking and ripples across the lifespan.

## 6 Acknowledgements

Special thanks to Michael Burke for apparatus construction.

