## Supplemental Figures 1-2, Table 1 for "Advanced Age Has Dissociable Effects on Hippocampal CA1 and CA3 Ripples"

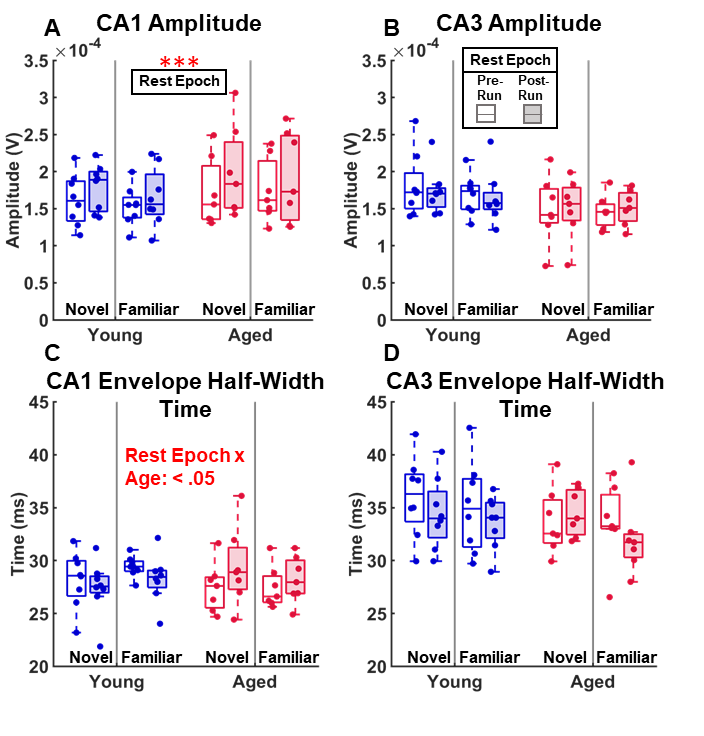

**Supplemental Figure 1:** Quantitative Measures for Young and Aged Ripples in CA1 and CA3 (continued): Maximum ripple amplitude **(A B)** and the time length of the ripple envelope at the half-amplitude **(C D)**. Individual data points are the median values for the representative metric for one rat (N(young)=8, N(aged)=7). Within each figure data is separated by age group, familiarity to the maze, and rest epoch. Metrics were tested for significance using a Mixed Model ANOVA (between: Age and Familiarity; within: Rest Epoch, p_critical = .05). CA1 amplitude had a significant effect with rest epoch (F _[1,26]_=16.46, p<.001) but this was not the same for CA3. CA1 ripple envelope half-amplitude width had a significant interaction between age and rest epoch (F _[1,26]_=5.02, p<.05) but CA3 did not.

| **Factor** | **CA1** | | **CA3** | |
| --- | --- | --- | --- | --- |
|  | **Amplitude** | **Envelope Half-Amplitude Time Width** | **Amplitude** | **Envelope Half-Amplitude Time Width** |
| **Age** | **F[1,26]=1.54 p=.23** | **F[1,26]=.07 p=.79** | **F[1,26]=3.59 p=.07** | **F[1,26]=1.39 p=.25** |
| **Day** | **F[1,26]=.16 p=.69** | **F[1,26]=.10 p=.76** | **F[1,26]=.22 p=.65** | **F[1,26]=.99 p=.33** |
| **Age x Day** | **F[1,26]=.10 p=.75** | **F[1,26]=1.27 p=.27** | **F[1,26]=.14 p=.71** | **F[1,26]=.02 p=.88** |
| **Rest Epoch** | **F[1,26]=16.46 p=.0004 ***** | **F[1,26]=.12 p=.73** | **F[1,26]=.12 p=.73** | **F[1,26]=2.45 p=.13** |
| **Age x Epoch** | **F[1,26]=.81 p=.38** | **F[1,26]=5.02 p=.03 *** | **F[1,26]=1.50 p=.23** | **F[1,26]=.92 p=.35** |
| **Day x Epoch** | **F[1,26]=.65 p=.43** | **F[1,26]=.56 p=.46** | **F[1,26]=.26 p=.61** | **F[1,26]=.81 p=.38** |
| **Age x Day x Epoch** | **F[1,26]=.05 p=.83** | **F[1,26]=.15 p=.70** | **F[1,26]=.0002 p=.99** | **F[1,26]=1.12 p=.30** |
| ***: p<.05 **: p<.01 ***:p<.001 #:p<.0001** | | | | |

**Supplemental Table 1: Statistical Results for Ripple Amplitude and Envelope Half-Amplitude Time Width**

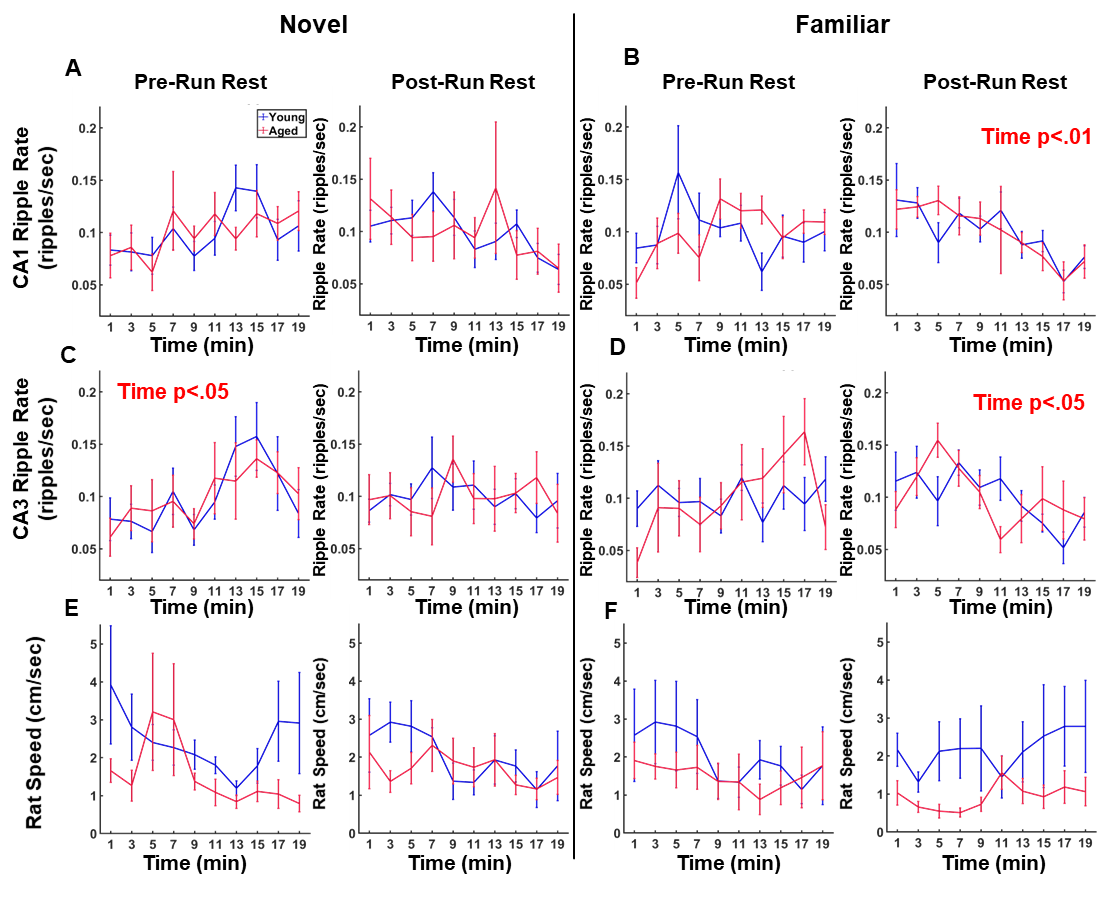

**Supplemental Figure 2: Ripple rate and rat speed calculated on two-minute blocks.** A Mixed Model ANOVA was calculated for each day/epoch across the different 2 minute time blocks (between: Age(2); within: Time Block(10)). Significance was only found for time block, but not age, in CA1 familiar post-run rest (F _[9,117]_=3.11, p<.01) **(B)**, CA3 novel pre-run rest (F _[9,117]_=2.12, p<.05) **(C)**, and familiar post-run rest (F _[9,117]_=1.99, p<.05) **(D)**. **(E-F)** Rat speed with the same time windows as ripple rate. There was no significant interactions for either age or time block for the Mixed Model ANOVA (between: Age, within: Time Block; F _[9,117]_<2.35, p>.15).
